# Potentials and limitations in the application of Convolutional Neural Networks for mosquito species identification using wing images

**DOI:** 10.1101/2025.01.29.635420

**Authors:** Kristopher Nolte, Jan Baumbach, Christian Lins, Jens Johann Georg Lohmann, Philip Kollmannsberger, Felix Gregor Sauer, Renke Lühken

## Abstract

**1.** This study addresses the pressing global health burden of mosquito-borne diseases by investigating the application of Convolutional Neural Networks (CNNs) for mosquito species identification using wing images. Conventional identification methods are hampered by the need for significant expertise and resources, while CNNs offer a promising alternative. Our research aimed to develop a reliable and applicable classification system that can be used under real-world conditions, with a focus on improving model adaptability to unencountered devices, mitigating dataset biases, and ensuring usability across different users without standardized protocols. **2.** We utilized a large, diverse dataset of mosquito wing images of 21 taxa and three image-capturing devices and an optimized preprocessing pipeline to standardize images and remove undesirable image features. **3.** The developed CNN models demonstrated high performance, with an average balanced accuracy of 98.3% and a macro F1-score of 97.6%, effectively distinguishing between the 21 mosquito taxa, including morphologically similar pairs. The preprocessing pipeline improved the model’s robustness, reducing performance drops on unfamiliar devices effectively. However, the study also highlights the persistence of inherent dataset biases, which the preprocessing steps could only partially mitigate. The classification system’s practical usability was demonstrated through a feasibility study, showing high inter-rater reliability. **4.** The results underscore the potential of the proposed workflow to enhance vector surveillance, especially in resource-constrained settings, and suggest its applicability to other winged insect species. The classification system developed in this study is available for public use, providing a valuable tool for vector surveillance and research, supporting efforts to mitigate the spread of mosquito-borne diseases.

## 1 Introduction

The global burden of mosquito-borne pathogens is of pressing concern. Diseases such as malaria, dengue fever, Rift Valley fever, or chikungunya cause significant morbidity and mortality worldwide, straining the public and veterinary health systems and impeding socio-economic development in affected areas (Franklinos et al., 2019). Furthermore, climate change and increased global trade contribute to the spread of mosquitoes and their associated pathogens (de Souza & Weaver, 2024).

The vector capacity of a mosquito species is a function of its ecology, behaviour, and vector competence, making accurate species identification a prerequisite for effective vector surveillance, research and control measurements (Kramer & Ciota, 2015). Conventional identification techniques, such as morphological analysis, are time-consuming and require specialist expertise, which is less and less available (Wilkerson et al., 2021). Alternative methods like molecular assays, can be prohibitively expensive and challenging to use in low-cost settings (Farlow et al., 2020). This concurs with the statement of the European Centre for Disease Prevention and Control, which underscores that a principal hurdle to the establishment of robust vector surveillance systems is the scarcity of trained experts and financial resources available (ECDC, 2021).

Within the context of insect species identification, machine learning (ML) driven methods have shown considerable promise, offering advancements that could automatize the identification process. Especially Convolutional Neural Networks (CNNs) have emerged as a powerful tool for identifying insects based on images, e.g. tsetse flies (Cannet et al., 2022), sandflies (Cannet et al., 2023) or bees (Buschbacher et al., 2020). For mosquitoes, several CNN models are presented in literature which are capable of classifying mosquito species based on images with varying scope and performance. For instance, Goodwin et al. (2021) achieved a remarkable 97% accuracy in identifying 17 mosquito species using standardized images of their bodies taken with a specifically designed imaging tower and Zhao et al. (2022) were even capable of distinguishing between species of the *Culex pipiens* complex, which cannot be morphologically differentiated by human eye. Although using images of whole mosquitoes may appear intuitive, leveraging wings for species classification provides distinct advantages. It has been demonstrated that using wing instead of body images not only improves classification performance but also reduces data demands for model training (Nolte, Sauer, et al., 2024). Wings are nearly two-dimensional, which makes the imaging process easily standardizable, as it does not require images from multiple angles. Moreover, wings and their vein patterns remain unaffected by the mosquito’s physiological status, such as being blood-fed or gravid (Li et al., 2024). Finally, wing landmark studies have shown that wings not only exhibit distinct interspecies differences but also intraspecies variations in wing vein patterns (Lorenz et al., 2017).

Traditional ML pipelines, while effective in many scenarios, frequently fall short in real-world deployment (Azulay & Weiss, 2019; Beery et al., 2018). ML development typically follows a standardized framework involving model specification, a training dataset, and an evaluation procedure that assumes identical distributions of training and deployment (D’Amour et al., 2022). However, this approach does not consider the potential for inherent biases and underspecifications within trained models. For instance, shortcut learning, as described by Geirhos et al. (2020), refers to a phenomenon where ML models learn to rely on simple, exploitable spurious correlations in the data (“shortcuts”) rather than the intended, more complex representations. For example, a model designed to identify cows in images might achieve high performance by leveraging the background context, such as associating the presence of grass with cows (Beery et al., 2018). These shortcuts often result in models that perform well on specific datasets but fail to generalize to new, unseen data (D’Amour et al., 2022; Geirhos et al., 2020). Another aspect of underspecifications is that CNN models are typically not specified to handle novel devices with varying imaging characteristics (D’Amour et al., 2022), a limitation also evident in insect (Fujisawa et al., 2023) and mosquito species identification (Nolte, Sauer, et al., 2024). Moreover, particularly in the context of image classification, deep learning methods are very data demanding. Thus, while the concept of a homogeneous dataset may seem ideal for a proof-of-concept study or in-house solutions, its practical implementation beyond such controlled environments is often unfeasible. These discrepancies between the training and deployment setting can lead to suboptimal performance in real-world conditions, and can thereby explain the often observed gap between proof-of-concept studies and practical applications in vector surveillance (Nolte, Sauer, et al., 2024).

Thus, there is a need for robust, open-access systems that can reliably identify mosquitoes under various image collection settings. Such an application holds the potential to facilitate vector monitoring, enabling a broader spectrum of applicants, including non-entomologists, to identify mosquitoes. In this study, we investigated the use of CNNs to classify mosquito species based on wing images, specifically addressing the challenges associated with deploying such models in real-world scenarios. Our primary goals were to 1) enhance model applicability to novel image-capturing devices i.e. devices which are not represented in the training set, 2) mitigate biases introduced by heterogeneous datasets collected with different devices and 3) ensure reliable use by different users without standardized image-capturing devices. To achieve these goals, we developed a classification system using a large, diverse dataset of mosquito wing images collected from multiple taxa and devices. This system incorporates an upstream image preprocessing pipeline to standardize images automatically and reduce the impact of undesirable features. We conducted a comprehensive evaluation of the system through four key approaches: in-distribution testing, simulation studies to assess the preprocessing pipeline’s effectiveness, explainable-AI methods to explore the models’ decision-making processes and feasibility study to evaluate the application’s practical use. Ultimately, we present a functional, openly available application for mosquito identification (GitHub Repository).

## 2 Materials and Methods

### 2.1 Dataset construction

A total of 14,888 images of female mosquito wings were sourced from an open-access mosquito wing dataset (Nolte, Agboli, et al., 2024). The dataset encompassed 8947 specimens from nine genera, *Aedes* (*Ae.*), *Anopheles* (*An.*), *Culex* (*Cx.*), *Culiseta* (*Cs.*), *Coquillettidia* (*Cq.*), *Armigeres* (*Ar.*), *Mansonia* (*Ms.*), *Urantaenia* (*Ur.*), *Toxorhynchites* (*Tx.*), and 72 species, species pairs, species groups and species complexes as defined by (Nolte, Agboli, et al., 2024). The images were often taken for both wings of the mosquito specimen, 48.2% of all specimens had both their wings imaged, and some wings were captured with multiple devices (11.2% of all wings).

To train the model, a specific label was assigned to each mosquito taxon. This label mostly refers to the mosquito species. However, the wing image dataset also includes specimens, which could not be identified at the species level as they are morphologically very similar or impossible to distinguish by classical taxonomy (e.g. species groups or species complexes). Therefore, the labels of the following taxa represent aggregated categories rather than discrete biological species: *Ae. annulipes* group, *Ae. cinereus*-*geminus* pair, *Ae. communis*–*punctor* pair, *An. claviger–petragnani* pair*, An. maculipennis* s.l., *Cs. morsitans*–*fumipennis* pair*, Cx. torrentium-pipiens* s.l. group, and *Cx. vishnui* group (Fig. 1). Taxa (N=34) represented with fewer than 80 images were summarised under the label “other” to ensure a robust training sample size per taxa label while still using all available wing images for model training. This threshold was based on previous experiments by Nolte et al. (2024). The labelling strategy resulted in 21 unique labels (Figure 1). Images were captured by two different stereomicroscopes (Olympus SZ61 in combination with the Olympus DP23 camera, Olympus, Tokyo, Japan and Leica M205 C, Leica Microsystems, Wetzlar, Germany) or a macro lens (Apexel-24XMH, Apexel, Shenzhen, China) attached to a smartphone (iPhone SE 3rd Generation, Apple Inc., Cupertino, USA).

**Figure 1:**
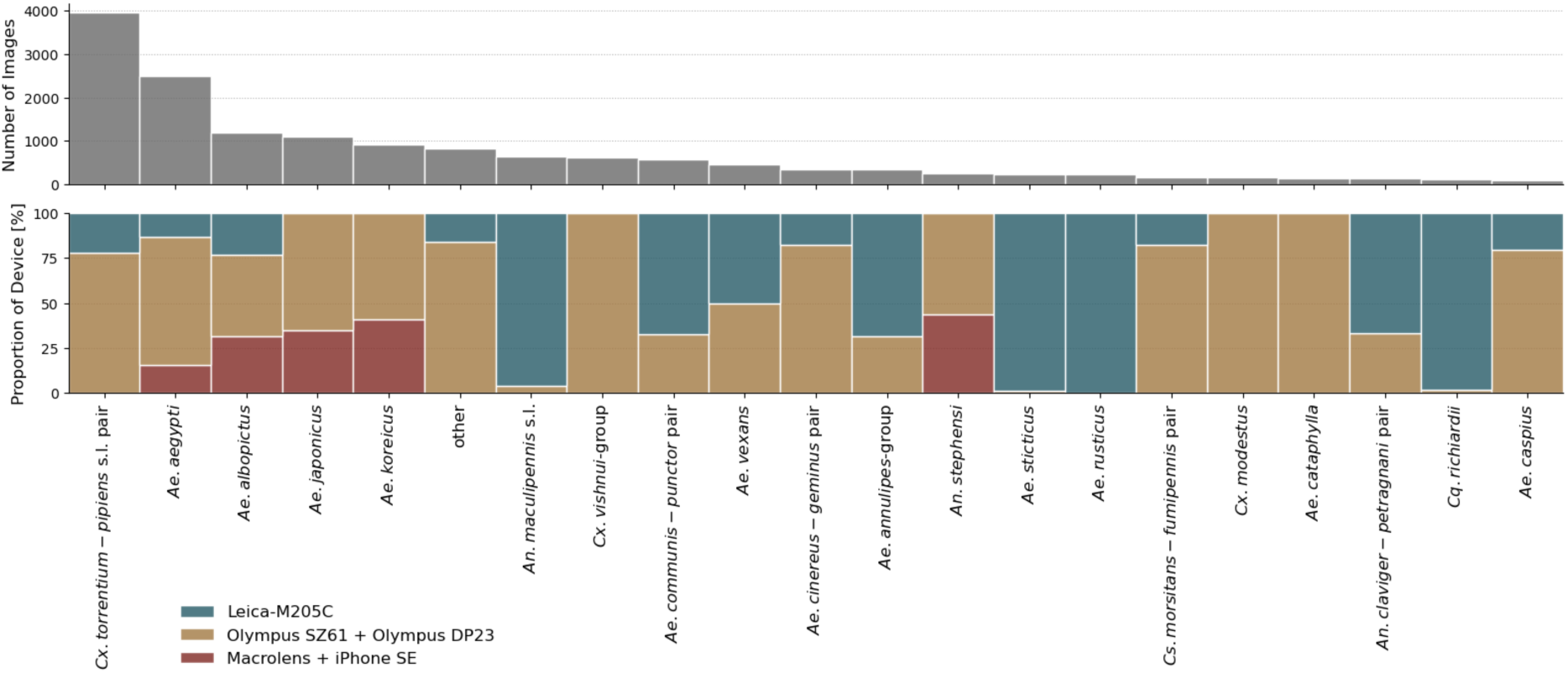
Number of images per taxa label (top) and proportion of image capture devices per label (bottom).

We implemented a six-fold cross-validation strategy to evaluate the performance of the CNN classification models. The dataset was randomly stratified into six folds, each containing approximately 2,500 images, ensuring proportional representation of the labels (Figure 2). Importantly, the partitioning was based on mosquito specimens rather than individual images. This approach was necessary because, in some of the underlying datasets, multiple images were taken of the same specimens. The images designated to the first fold were reserved for validating and tuning the model’s hyperparameters. The other five folds were later used for training and testing. In each training iteration, images designated to one-fold were designated as the test set while the remaining images were used for model training. Comprehensive details about the images, including their labels, capture devices, and sources, are provided on the publications’ GitHub page (<u>GitHub Re-pository</u>).

**Figure 2:**
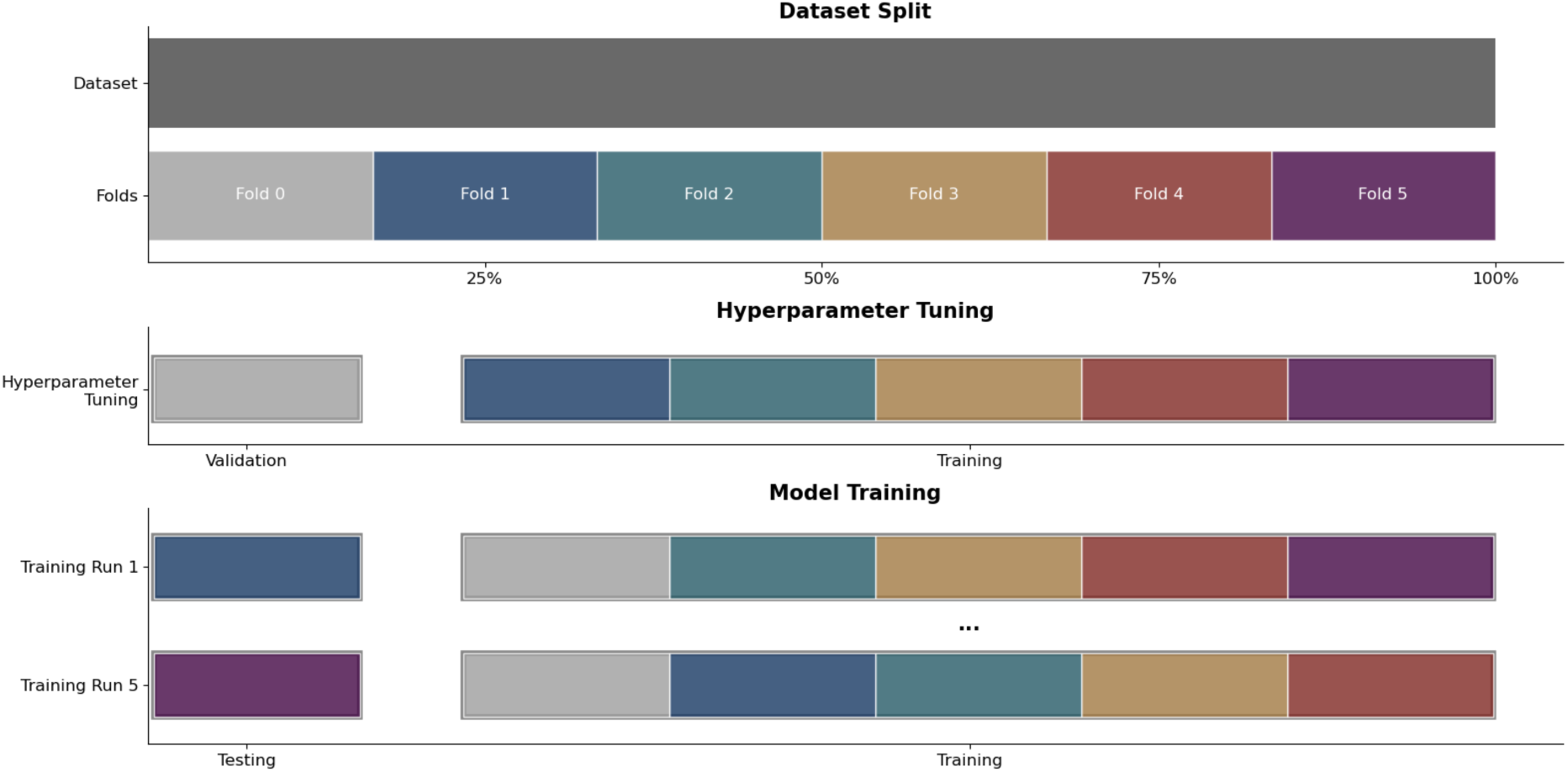
Plot illustrating the division of the image dataset into six folds (Top). First, the model hyperparameters were optimized by utilizing the first fold (fold 0) as the validation set (Middle). The bottom sections depict the training strategy, showcasing cross-testing across folds to generate five unique dataset compositions, with one fold designated as a testing set in each training run.

### 2.2 Preprocessing pipeline and augmentation

We developed an image preprocessing pipeline, aiming to reduce inherent biases in our heterogenous dataset and improve the model’s applicability to images captured by novel devices (Figure 3). The pipeline aims to eliminate undesired features such as varying lighting conditions, background, and colour variations. In the first step, the images are loaded (Figure 3, step: A) and hereafter a U-Net model from the Rembg Python library is employed in the second step to create a segmentation mask of the wing and background to remove the latter (step: B, <u>github.com/danielgatis/rembg</u>). Next, in the third step, the image is augmented (step: C). To implement this, we utilized augmentation functions from the Albumentations library to construct an augmentation pipeline consisting of 13 layers: ISONoise, PlanckianJitter, ImageCompression, Defocus, RandomGamma, MotionBlur, Downscale, ColorJitter, ChannelDropout, MultiplicativeNoise, RandomZoomOut, RandomHorizontalFlip, and RandomRotation (Buslaev et al., 2018). During training the augmentation is used to increase the variance of the training set, during deployment the augmentation is used for test-time augmentation, only during testing this step is omitted. In the fourth step, orientation of the wings is horizontally aligned by using region properties measured from the segmentation mask of the wing (step: D), implemented with sci-kit-image (Walt et al., 2014). This alignment adjusts the horizontal orientation of wings across different images, reducing variability and aiding consistent feature extraction. To reduce the impact of colour variations, which are generally not very informative for traditional mosquito species identification by wings (Sauer et al., 2024), the images are converted to greyscale (step E). This conversion simplifies the image while retaining essential structural information, improving the robustness of the model to varying colour changes due to different imaging settings. Contrast Limited Adaptive Histogram Equalisation (CLAHE) with a high clip limit (0.5) is applied (step F) before median filtering (Pizer et al., 1984). CLAHE enhances image contrast and improves the visibility of the wing vein pattern. Median filtering smooths out noise and further enhances the clarity of the wing details. Lastly, images are cropped and resized to a resolution of Y:192 and X:384 to ensure they are suitable for input into the CNN model (step G). This size is chosen because it keeps enough detail for the model to recognize important features while also reducing the computational effort required for training.

**Figure 3:**
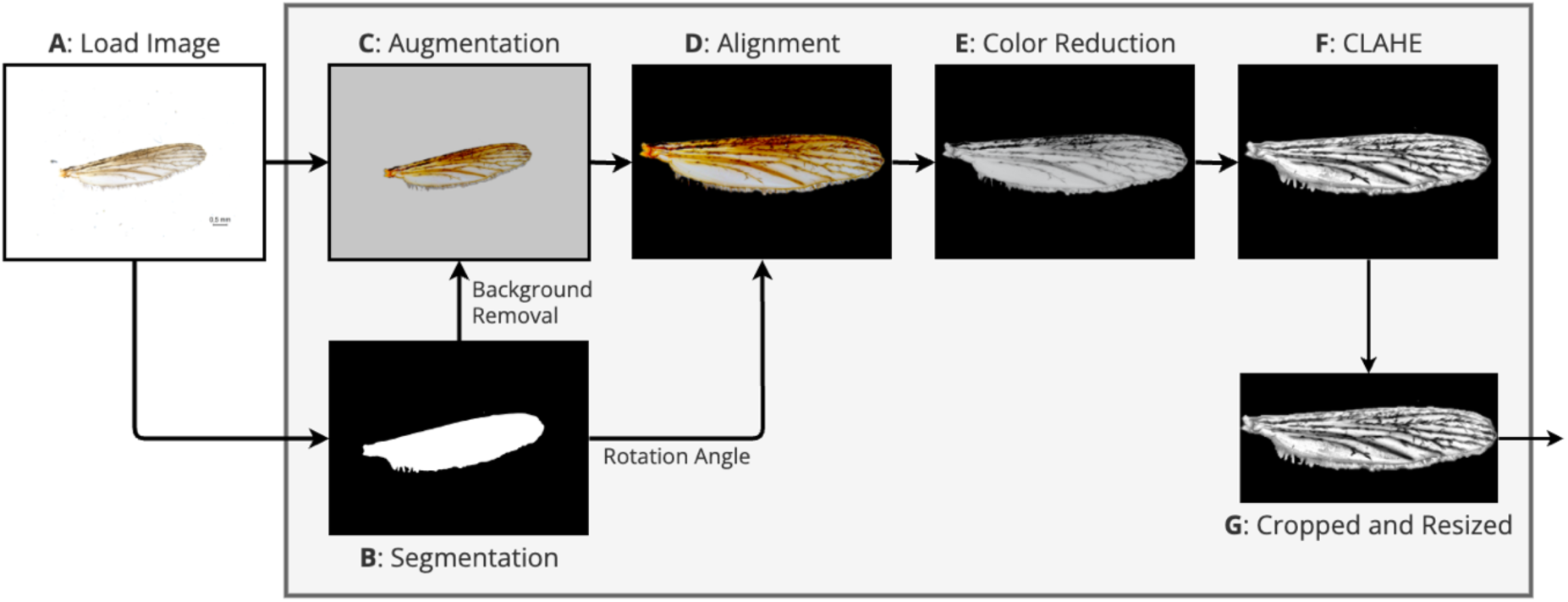
Schematic representation of the image preprocessing pipeline. The process starts with the loading of the original wing image (A). Background removal is performed to isolate the wing using a segmentation mask (B). The image is augmented by an augmentation pipeline (C). The augmented wing is then rotated to a standardized horizontal orientation based on the calculated rotation angle (D). The image is converted to greyscale to reduce colour variability (E). CLAHE is applied to enhance contrast and highlight important features (F). Finally, the processed image is resized to a uniform dimension, making it suitable for input into the CNN model (G).

### 2.3 Classifier Training

A training setup was developed, encompassing the pipeline and training procedure using the Python programming language and the PyTorch library. The Python scripts and a list of all used libraries and their versions can be found on the publications GitHub page (GitHub Repository).

The issue of imbalanced data distribution is addressed by resampling a uniform distribution from the training dataset in each training epoch. The training process was facilitated by using pre-trained CNNs and the fine-tuning learning strategy. The EfficientNetB0 model architecture without the original classification head, pre-trained on the *ImageNet* dataset, was utilized as a feature extractor (Deng et al., 2009; Tan & Le, 2019). Additional layers, namely a Dropout and a Dense Layer, were appended for classification.

Hyperparameters such as learning rate, batch size, and image size were systematically optimized through experimental iterations and evaluation of the validation set. For model training, categorical crossentropy emerged as a loss function for the classification task. The Adam optimizer with weight decay was employed to optimize the learned parameters during model training (Loshchilov & Hutter, 2019). A complete list of hyperparameters resulting in the final models can be found in the supporting information (supporting information: Hyperparameter). After the hyperparameters were determined, five models were trained and tested on five folds to acknowledge the stochastic nature of ML models (Figure 2). The mean balanced accuracy and macro F1-score of the five models are reported with a 95% confidence interval (CI95%) on the testing set.

### 2.4 Robustness experiments

To investigate the efficacy of our preprocessing pipeline in reducing model applicability to unfamiliar devices and biased data, we designed two experiments (“novel-device” and “bias”-experiment) using the systematically collected mosquito wing dataset from Nolte et al. (2024). The dataset contains 3,027 wing images of 793 female specimens of *Ae. aegypti, Ae. albopictus, Ae. koreicus*, and *Ae. japonicus* each captured with both, a microscope (Olympus SZ61 in combination with the Olympus DP23 camera, Olympus, Tokyo, Japan) and a macro lens (Apexel-24XMH, Apexel, Shenzhen, China) attached to a smartphone (iPhone SE 3rd Generation, Apple Inc., Cupertino, USA). This dataset was divided into five folds using the same protocol described in Section 3.1 (Figure 2), except that no fold was designated for validation since all hyperparameters were kept constant. The only adjustment was a reduction in training epochs from 32 to 12 and the freezing of the first half of layers of the model to account for the smaller dataset size and prevent overfitting. All images of the training data underwent four levels of increasing preprocessing based on intermediate steps of the preprocessing pipeline: A = original images only resized with padding (Fig. 3.A), C = with augmentation (Fig. 3.B), D = with background removed and wings horizontally aligned, these images were cropped and resized without padding (Fig. 3.D) and G = images processed by the complete preprocessing pipeline (Fig. 3.G). In the first experiment, i.e. the novel-device experiment, we aimed to evaluate the applicability of the CNN model to images from previously unencountered devices, examining its performance across the four levels of preprocessing. Using the wing images of the four *Aedes* species, we constructed two separate datasets: one consisting solely of smartphone-captured images and the other consisting solely of microscope-captured images. In two training sessions, we trained five models on either smartphone-captured images or microscope-captured images, as described in section 2.3. Subsequently, the performance of these trained models was tested on the respective testing fold, containing only images captured with the same device. In addition, the model was also tested on all images of the dataset which were captured using the device not present in the training or testing data.

The second experiment, i.e. the bias experiment, was designed to evaluate the reduction of model bias through the four levels of preprocessing, using a dataset with inherent biases. In this biased dataset, only smartphone images of the two species *Ae. albopictus* and *Ae. japonicus* and only microscope images of the two species *Ae. aegypti* and *Ae. koreicus* were included. The goal was to increase the likeliness of the models to utilise the spurious correlation of imaging characteristics with the species label. We split the dataset into 5 folds and trained five models on the different dataset compositions. During testing, we not only tested on the testing folds of the biased dataset but also on the remaining images not present in the biased dataset which had the inverse species–device combinations e.g. *Ae. aegypti* is here represented by images captured by microscope. This approach allowed us to assess the degree to which the different levels of preprocessing reduced the model’s bias toward using device-specific imaging characteristics for classification.

### 2.5 Model exploration

We employed Gradient-based Class Activation Mapping (GradCam) to gain insights into the decision-making process of the CNN models. GradCam calculates the gradients of the target class for the final convolutional layer of the CNN, highlighting regions with high gradients to produce a heatmap visualization of the most discriminative image regions (Selvaraju et al., 2020). Recognizing the potential challenges in interpreting Grad-Cam heatmaps (Chattopadhay et al., 2018), we applied GradCam to multiple images in the test dataset, averaging the resulting activation maps and images. This approach was facilitated by our preprocessing pipeline, which ensured that all images were consistently aligned. For this analysis, we utilized the best-performing model and mosquito wing images from the Sauer et al. (2024) study, as this dataset provided uniformly oriented images of the right wing. We focused on six taxa labels from the dataset as these provided the most images: *Ae. cinereus-geminus*-pair, *Ae. communis-punctor*-pair, *Cq. richiardii*, *Ae. rusticus*, *Ae. sticticus*, and *Ae. vexans*. For each taxon, we randomly selected 12 images from the testing fold, applied GradCam to these images, and subsequently averaged both, the GradCam activation maps and the original images, to create a composite visualization of the most discriminative regions for each taxon.

To further investigate the decision-making process of the CNN models, we applied Uniform Manifold Approximation and Projection (UMAP) to the feature maps generated by the best-performing CNN model. UMAP is a dimensionality reduction technique that enables visualization of high-dimensional data in a lower-dimensional space while preserving the underlying structure and relationships (McInnes et al., 2018). By applying UMAP to the feature maps, we aimed to uncover patterns and relationships within the data that may not be apparent in the original high-dimensional space.

### 2.6 Feasibility study

A feasibility study was conducted to assess the applicability of the CNN model under real-world conditions. Four participants were tasked to independently capture images of mosquito wings of 10 taxa labels, comprising a total of 78 wings: *Ae. aegypti* (8 images), *Ae. albopictus* (8), *Ae. japonicus* (8), *Ae. koreicus* (8), *Ae. vexans* (8), *Cx. pipiens* s.l./*Cx. torrentium* (8), *Cx. tritaeniorhynchus* (8), *An. maculipennis* s.l. (6), *An. stephensi* (8), *Cq. richiardii* (8). Additionally, we included 8 wings belonging to the “other” label (*An. coustani*-group). Participants utilized a stereomicroscope (Olympus SZ61, Olympus, Tokyo, Japan) equipped with a camera (Olympus DP23, Olympus, Tokyo, Japan). Subsequently, all captured images underwent processing through the image preprocessing pipeline. The processed images were then classified with all 5 models.

### 2.7 Metrics

The accuracy and F1-score were used as performance metrics (Table 1). To adapt the performance metrics for the imbalanced multilabel classification, we used balanced accuracy and macro F1 score. We assessed the interrater reliability, by utilizing the Fleiss’ Kappa, which measures the agreement between multiple raters when assigning categorical ratings to a fixed number of items (Shrout & Fleiss, 1979). To calculate the metrics, we utilised the Python libraries scikit-learn and statsmodels (Pedregosa et al., 2011; Seabold & Perktold, 2010).

**Table 1:** Metrics and their definition which were used to assess the performance of the classification system. TP: True positives, TN: True negatives, FP: False positives, FN: False negatives.

| Metric | Equation |
| --- | --- |
| Accuracy | $\frac{TP + TN}{TP + FP + TN + FN}$ |
| Precision | $\frac{TP}{TP + FP}$ |
| Recall | $\frac{TP}{TP + FN}$ |
| F1 Score | $2 * \frac{Precision * Recall}{Precision + Recall}$ |

### 2.8 Model deployment

To make the image preprocessing pipeline and the model accessible, we developed a user-friendly application using the Python framework Flask (Grinberg, 2018). This application provides a web service that allows users to upload images of mosquito wings, which are processed through the defined imaging pipeline, augmented multiple times (test-time augmentation) and then analyzed by the best-performing CNN model for prediction. The prediction from four augmented versions of the wings is then averaged and calibrated to produce an output between 0 and 1 representing the uncertainty of the model (Guo et al., 2017). Upon uploading images, users receive a display of the predictions and a CSV file containing the models’ averaged, calibrated predictions. To ensure consistency across different deployment environments, we employed Docker to containerize the entire classification system, encapsulating all dependencies and configurations. This approach simplifies installation and enables seamless execution regardless of the operating system or environment. The application, along with installation instructions, is available on GitHub and can be downloaded and run on a computer (GitHub Repository). Additionally, the application is hosted on a website (https://balrog.bnitm.de). However, due to legal constraints, e.g. uploading of copyright-protected images, access is currently password protected. Interested users are encouraged to contact the authors to request access credentials.

## 3 Results

### 3.1 Classifier Performance

Following the selection of the final hyperparameters, five models were trained on the five training configurations and subsequently evaluated on the remaining images of the respective testing fold. The CNN models exhibited high performance, achieving an average balanced accuracy of 98.2% (CI95%: 97.8-98.6) on the testing sets (Figure 4). The average macro F1-Score was 97.6% (CI95%: 96.9-98.4). The model that has been tested on fold 2 has shown the best performance in terms of balanced accuracy with 98.6%. In total, summarizing all testing folds, 214 of 12323 images (1.7%) were misclassified. Among the misclassified images, 26 (12%) underwent undesirable preprocessing, while 30 wings (14%) exhibited damaged features. The classification accuracy varied only a little across different taxa, with 2 out of 21 labels achieving flawless classification (100%), and 19 labels attaining accuracy levels exceeding 95%. The CNN models demonstrated lower average accuracy (≤ 95%) only for the “other” taxon label.

**Figure 4:**
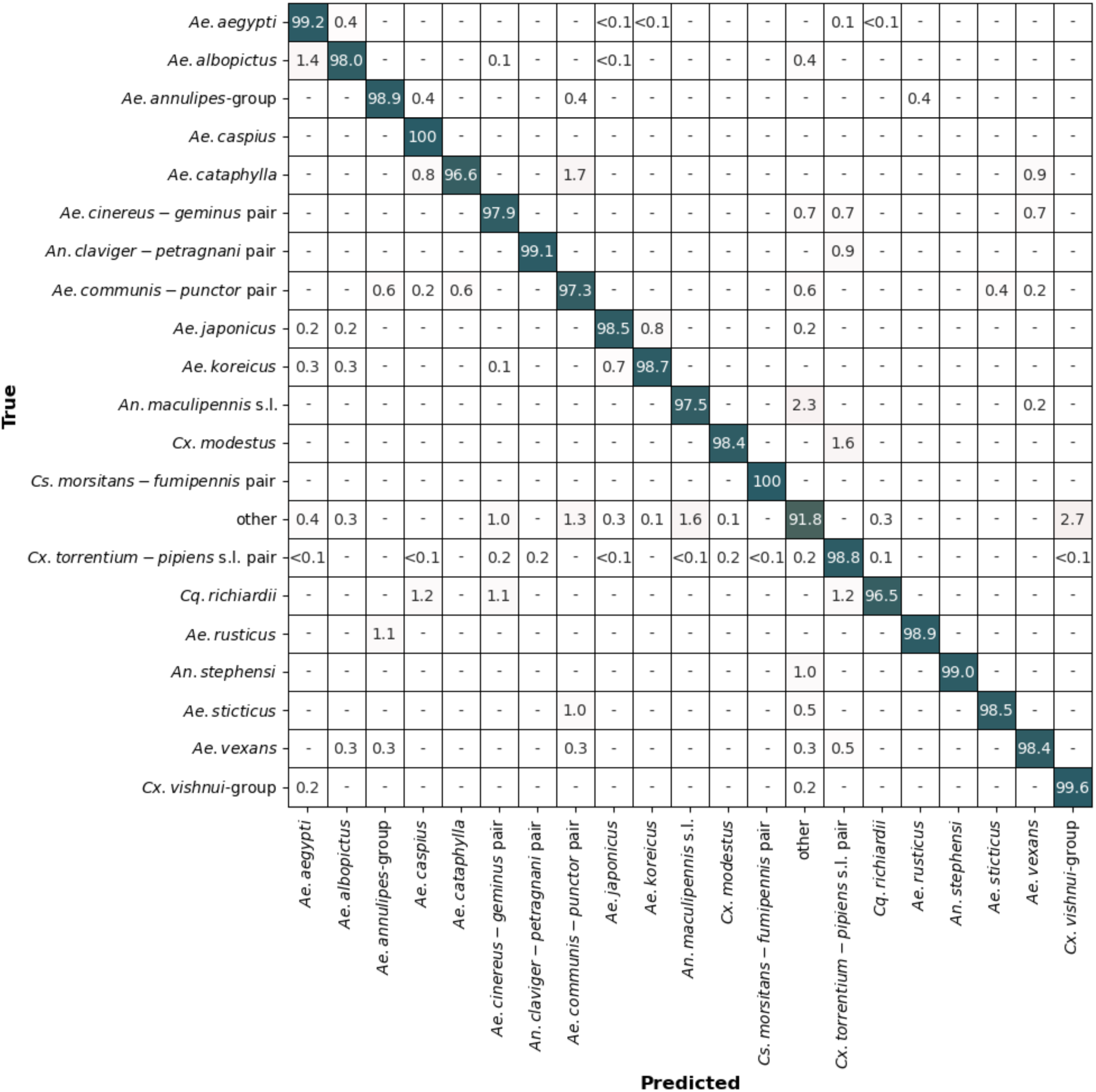
Confusion matrix showing the average accuracy of the five CNN models in classifying mosquito taxa based on wing images. Each cell represents the proportion of predictions made for each true taxon (%).

### 3.2 Robustness experiments

In the novel-device experiment, we assessed the models’ applicability to images captured with unfamiliar devices by testing the performance under different levels of preprocessing. The models’ performance on images from a new device improved with higher levels of preprocessing, while their performance on the original training device remained stable (Figure 5.1a and 5.1b). The highest performance on a novel device is achieved by the full processing method. It resulted in an average accuracy of 90.0 (CI95%: 87.2-92.9) and 94.5% (CI95%: 92.2%-96.7%) when trained only on microscope or macro lens images, respectively. This means a performance drop of 7.8% for the microscope-trained models and 2.1% for the macro-lens-trained models. In contrast, with minimal preprocessing (only resizing), the performance drop was much larger: 75.2% for the microscope-trained models and 62.4% for the macro-lens-trained models.

**Figure 5:**
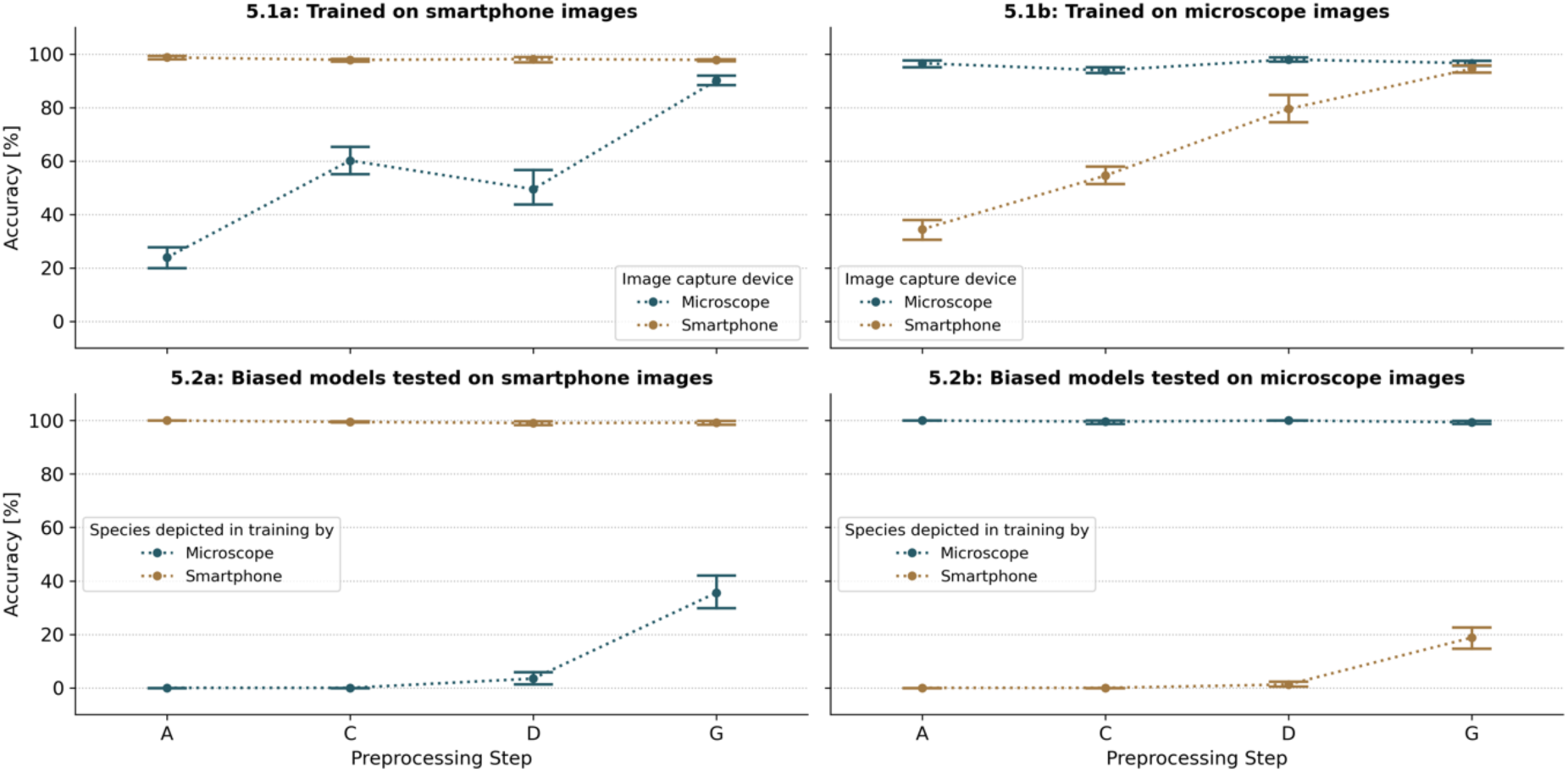
Impact of preprocessing (A = original images only resized with padding (Fig. 3.A), level C = with augmentation (Fig. 3.C), C = with background removed and wings horizontally aligned, these images were cropped and resized without padding (Fig. 3.C) and G = images processed by the complete preprocessing pipeline) on model performance across different training conditions. Panel 5.1a and 5.1b display results from the novel-device experiment, Panel 5.2a and 5.2b from the bias experiment. Panels 5.1a and 5.1b show the accuracy of models trained on macro lens images (5.1a) and microscope images (5.1b) tested on both, the original device and a new device. Panels 5.2a and 5.2b show the accuracy of models tested on new species-device combinations, with training data biased towards one device. Panel 5.2a displays results for macro lens images, while Panel 5.2b displays results for microscope images.

In the second experiment, we introduced bias into the training dataset by using exclusively smartphone images for two mosquito species and microscope images for the other two species. When tested on the same species device combination as in the training dataset we observed a nearly perfect classification performance (>99%) on all preprocessing levels. Yet, as anticipated, the models performed poorly when tested on novel combinations of species and imaging devices. Without preprocessing, the models failed to correctly predict any images with a novel species-device combination (Figure 5.2a and 5.2b). However, increasing the level of preprocessing resulted in marginal improvements in model performance and a substantial boost was observed when full preprocessing was applied. For images captured with a smartphone, the models achieved an average accuracy of 35.7% (CI95%: 25.1%-46.3%) on novel species-device combinations when full preprocessing was applied (Figure 5.2a). A similar, but less pronounced, improvement was observed with microscope images, where accuracy increased by 18.7% (CI95%: 12.3%-25.2%) under full preprocessing (Figure 5.2b).

### 3.3 Model evaluation

GradCam analysis was performed to provide insights into the decision process of the CNN models for six species. When wing images were averaged, the resulting depictions clearly show individual veins, demonstrating the effectiveness of the preprocessing pipeline in consistently aligning the wings (Figure 6). The Grad-Cam images revealed variations in the importance of regions used for classification across different species. Across all species, the visualizations consistently emphasize structural patterns near the central regions of the wing. The primary regions of importance appear to correspond to intricate vein arrangements and interveinal membrane structures. For all taxa examined, the highest activation regions are observed along the medial section of the wing, where vein branching is most complex, while peripheral areas (distal tips and edges) are less prominently highlighted. Notably, species such as *Cq. richiardii* and *Ae. cinereus*-*geminus*-pair exhibit broader and more diffuse activation patterns, compared to the tighter focal regions observed for taxa like *Ae. rusticus* and *Ae. sticticus* (Figure 6).

**Figure 6:**
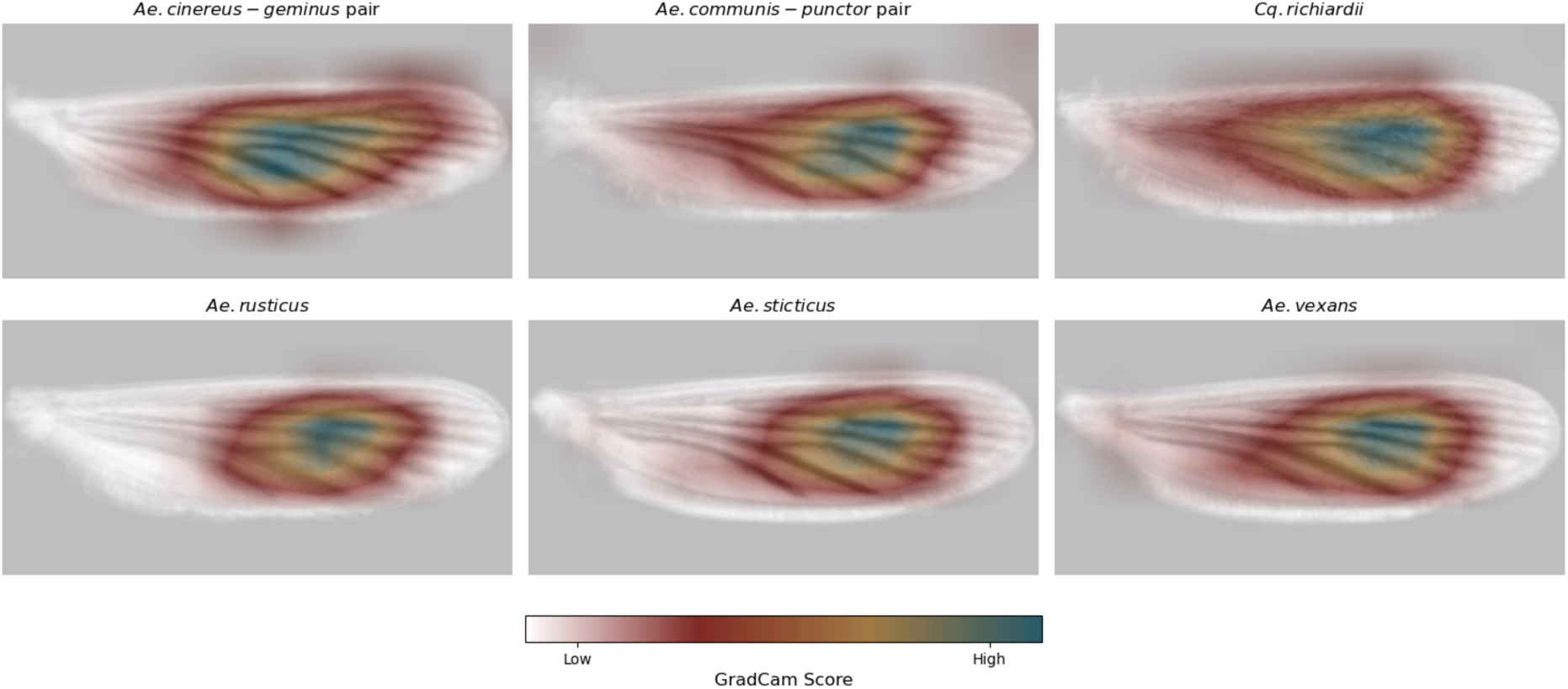
Averaged GradCam heatmaps generated from 12 wing images for six mosquito taxa labels. The heatmaps highlight the regions of the wings that the CNN model focuses on for classification, with blue areas indicating higher importance and yellow to red areas indicating lower importance while white areas are of no importance.

To explore the distribution of taxa within the feature space, UMAP analysis was performed on the feature maps of the testing fold 2 for the corresponding CNN model. The results reveal distinct clustering patterns, with most taxa forming clearly defined and well-separated clusters (Figure 7). Instances of proximity between clusters were observed, particularly among morphologically similar taxa, e.g. the close positioning of *Ae. aegypti* and *Ae. albopictus* or *Ae. koreicus* and *Ae. japonicus*. These observations align with the morphological challenges inherent in distinguishing these taxa, as their wing features exhibit significant similarity. Additionally, images belonging to the “other” species label form a broad, less cohesive cluster, reflecting the heterogeneous nature of this category. The UMAP analysis can also illustrate the influence of imaging devices on feature space organization (Figure 7) where no distinct clusters corresponding to different capture devices are observed.

**Figure 7:**
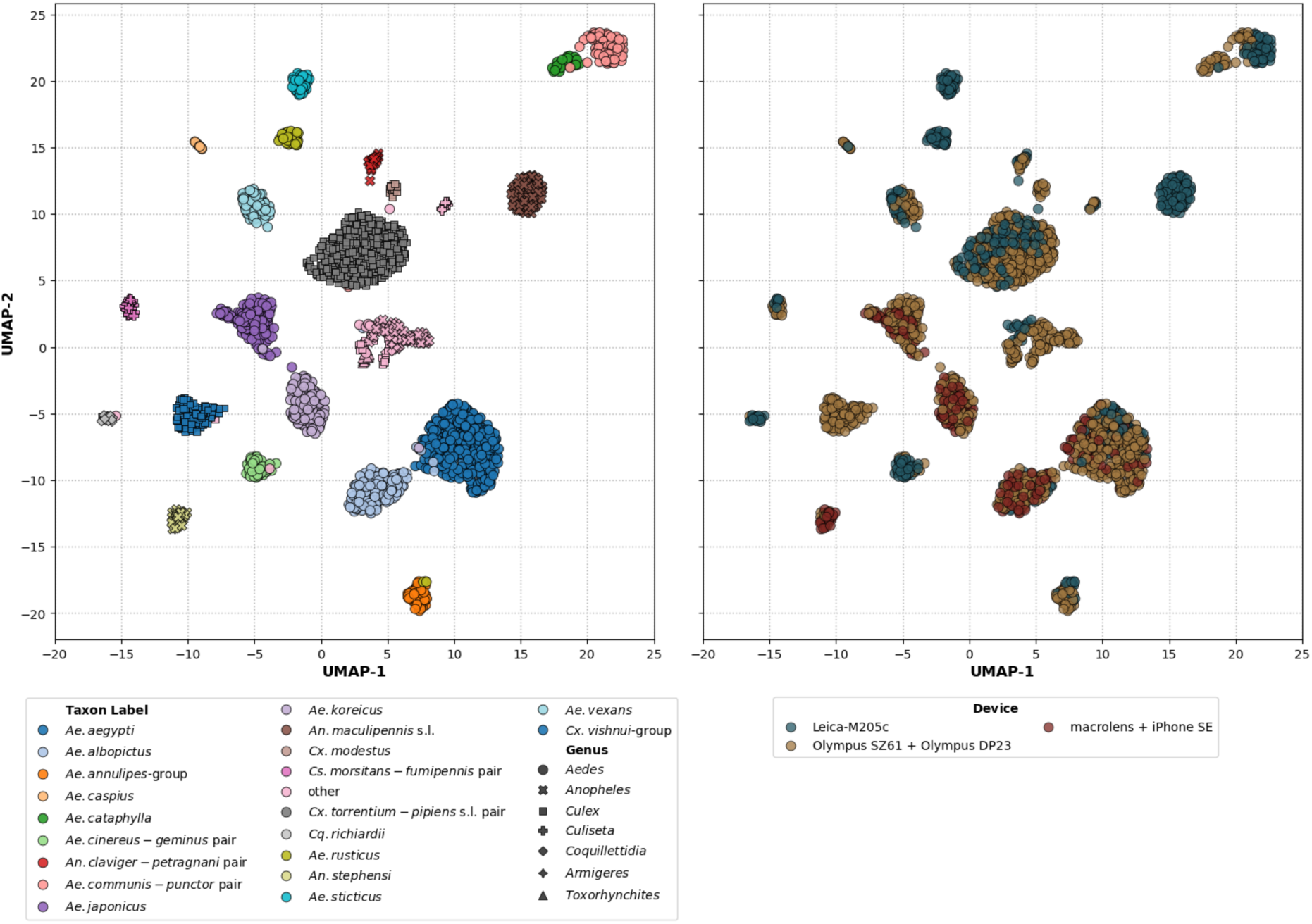
UMAP on the feature maps of the training fold 1 for the corresponding CNN model with the mosquito taxa colour-coded (left) and imaging devices-coloured (right).

### 3.4 Feasibility study

The feasibility study aimed to evaluate the applicability of the developed mosquito classification system in a real-world scenario. During the study, four participants were tasked to apply the developed models for 86 samples of mosquito wings which were neither represented in the training nor testing datasets. The five models demonstrated an average balanced accuracy of 96.4% (CI95%: 95.9-97.0). The average balanced accuracy is 1.4% lower than the expected 97.8% based on the average taxa label accuracy on the testing set. The worst performance was observed for *Cq. richiardii* which was only classified correctly with 72.5% accuracy. Most misclassifications were concentrated in two samples, which were consistently misclassified over all participants and models. There was no apparent reason for the complete misclassification of these samples. Despite these errors, the model achieved consistent predictions regardless of the participant. The Fleiss’ Kappa coefficient, calculated for all predictions divided by participants, was 0.97, indicating a very high level of agreement of the predictions.

## 4 Discussion

This study investigated the application of CNNs to identify mosquito species based on wing images for vector surveillance and research. The objective of the study was not only to develop a reliable CNN model for the classification of mosquito wings but also to ensure that the model was applicable under real-world settings, including compatibility with different imaging devices and users. The developed classification models demonstrated a high performance both, measured in average balanced accuracy (98.2%) and average macro F1-score (97.6%). Notably, the models showed high performance across 21 taxa labels even with morphologically similar pairs, e.g. *Ae. aegypti* and *Ae. albopictus* or *Ae. koreicus* and *Ae. japonicus*. The GradCam images indicate that the models are using reasonable features for wing classification. Additionally, the UMAP visualisation of a feature map shows that the model is grouping the different species labels into distinct clusters, indicating that the model has learned similar features from the images of the same species label. The most frequently misclassified species labels were *Cq. richiardii*, which had one of the fewest images in the dataset (N=104), and the ‘other’ label, which included a diverse composition of less frequently represented taxa.

Compared to previous studies using neural networks for wing-based mosquito identification, our models demonstrated better performance both in the number of taxa classified and overall accuracy relative to the findings reported in the existing literature (Cannet, Simon-Chane et al., 2023; Sauer et al., 2024). Even when compared to species identification through morphology the classification presents satisfying performance, e.g. Rahola et al. (2022) reported an average accuracy of just 81.5% for species identification based on responses from 51 participants provided with detailed images of mosquito samples. This difference highlights the potential of CNN-based identification methods to surpass the limitations of traditional morphology-based approaches, which heavily depend on the expertise of the observer. However, it is important to note that our model’s performance is inherently restricted to the species it was trained on, and its utility in broader applications will depend on the availability of training data for additional species.

The image preprocessing pipeline proposed in this study enhances the model’s applicability to new devices that were not included in the training data. In our novel-device experiment, we found that increasing levels of preprocessing improved accuracy on new capture devices from 23-34% without preprocessing to 90-93% with full preprocessing, which is comparable to the model’s accuracy on images from devices included in the training data. At the same time, preprocessing does not compromise the model’s accuracy on the images that were captured with familiar devices, indicating that no essential features for classification are lost during preprocessing. While data augmentation is often recommended to improve the generalizability of models to new settings (Ilse et al., 2021), our experiments revealed that it only partially enhances adaptability to new devices and does not improve robustness to bias. The argument for preprocessing is further supported by Fujisawa et al. (2023), who identified the lack of standardization as a key factor contributing to reduced accuracy when applying CNN models beyond their original training distributions.

However, the bias reduction experiment still highlights some limitations of the preprocessing pipeline as it could only partially mitigate the effects of the highly skewed training dataset used in the experiments with a strongly biased distribution between two very different image capture methods. These models only achieved an average balanced accuracy of 35.8% on novel species-device combinations. Consequently, one must exercise caution when adding new species to the model’s training data as device-specific variations in image characteristics may inadvertently introduce biases that the current preprocessing pipeline cannot fully mitigate. We recommend adhering to a threshold of at least 100 images per species to ensure reliable training, consistent with findings from Nolte et al. (2024) and our observations for *Cq. richiardii*. For rare or underrepresented species, grouping them under the “other” label offers a practical alternative. This strategy would allow the model to reject the predictions, thereby reducing false positives while preserving overall reliability. In addition, in future research, the issue of a device-specific bias could for example be addressed by utilising contrastive learning, which trains the model to learn an embedding space where similar data points, i.e. species labels, are close together and dissimilar points are far apart (Zhang et al., 2022).

Wings, as opposed to bodies, appear to be more reliable for this classification task. We applied heavy preprocessing methods, which most likely would have destroyed important features for automatic full mosquito body classification, e.g. omitting legs during background removal. While the removal of wings is an additional work step, wing vein patterns themself present as robust features, which are not associated with colour or texture, allowing to reduce the presence of undesirable features in the images without reducing the model performance. Furthermore, the depiction of whole mosquitoes can vary significantly due to factors such as age, physiological status (e.g. blood-fed or gravid), storage conditions, and image angel introducing additional variability that complicates the classification process (Couret et al., 2020; Li et al., 2024). Thus, the relatively stable and distinct structures of the wings probably provide a more reliable basis for preprocessing and subsequent classification.

While the presented classification model demonstrates strong performance, it is currently restricted to a limited selection of mosquito taxa primarily native to Europe (Robert et al., 2019). This limitation introduces uncertainty regarding the model’s ability to identify previously unencountered species. Specifically, the “other” label, designed as a catch-all category, is unlikely to adequately detect entirely new species due to the relatively narrow representation of taxa within the “other” label distribution (Hendrycks & Gimpel, 2018). Additionally, the wing shape and size within a species can slightly vary across mosquito populations from different regions, influenced by environmental and genetic factors (Francuski et al., 2016; Gómez et al., 2014; Hounkanrin et al., 2023). Similarly, (Fujisawa et al., 2023) demonstrated that CNNs trained on globally distributed images of beetles (*Coleoptera*) exhibit reduced performance when applied to a new, local dataset. While the extent to which this effect applies to mosquito wings is unknown, we anticipate that the models presented here may also demonstrate a performance decline when tested on unfamiliar local populations. Together, these factors underscore the importance of validating the classification system under novel conditions and, when necessary, tailoring it to specific ecological contexts.

The feasibility study supports the classification system’s usability under real-world conditions, demonstrating high inter-rater reliability with a Fleiss’ kappa of 0.97 and a similar balanced accuracy compared to the testing performance. The ability of the preprocessing pipeline to enhance the model’s adaptability to previously unencountered devices and different users highlights its potential for application in vector surveillance and research. The preprocessing methods reduce the need for stringent standardization protocols or specific image devices and address the needs of current vector surveillance and research, targeting both major hurdles: lack of funding and trained personnel (ECDC, 2021; Farlow et al., 2020). The here presented method only requires tweezers, and a macro-lens attached to a smartphone and can be conducted by a nonexpert user. This effectively reduces the barriers to integrating this method into vector surveillance efforts and could facilitate its use in resource-constrained settings (Fournet et al., 2018). In the future, the model for mosquito wing classification can be extended to other dipteran insects of interest, e.g. biting midges or sandflies. These insects also exhibit distinct wing patterns that can be captured and analysed using similar preprocessing and classification techniques (Cannet et al., 2022; Sereno et al., 2023).

## 5 Additional Notes

### Authors’ contributions

KN, FGS, PK, CL and RL conceived the ideas and designed methodology; KN, FGS and RL collected the data; KN, JJGL developed models and software; KN, FGS, JJGL, JB, PK, and RL analyzed the data; KN, FGS and RL led the writing of the manuscript. All authors contributed critically to the drafts and gave final approval for publication.

### Data availability

Images used in this study are freely available and open access to be found on: https://doi.org/10.6019/S-BIAD1478

Scripts which were used to train models are available on the project git page: https://anonymous.4open.science/r/MosquitoWingClassifier_publication-3780

Downloadable and installable docker application can be accessed on the applications’ git page: https://anonymous.4open.science/r/MosquitoWingClassifier-7533

The web-hosted application can be accessed using this address, please note that a password will be required, please contact the authors regarding access: https://balrog.bnitm.de

## 7 Supporting information

### 7.1 Glossary

CI95%: 95% confidence interval
CNN: Convolutional Neural Network
CLAHE: Contrast Limited Adaptive Histogram Equalisation
GradCam: Gradient-weighted Class Activation Mapping
ML: Machine Learning
UMAP: Uniform Manifold Approximation and Projection

### 7.2 Hyperparameters

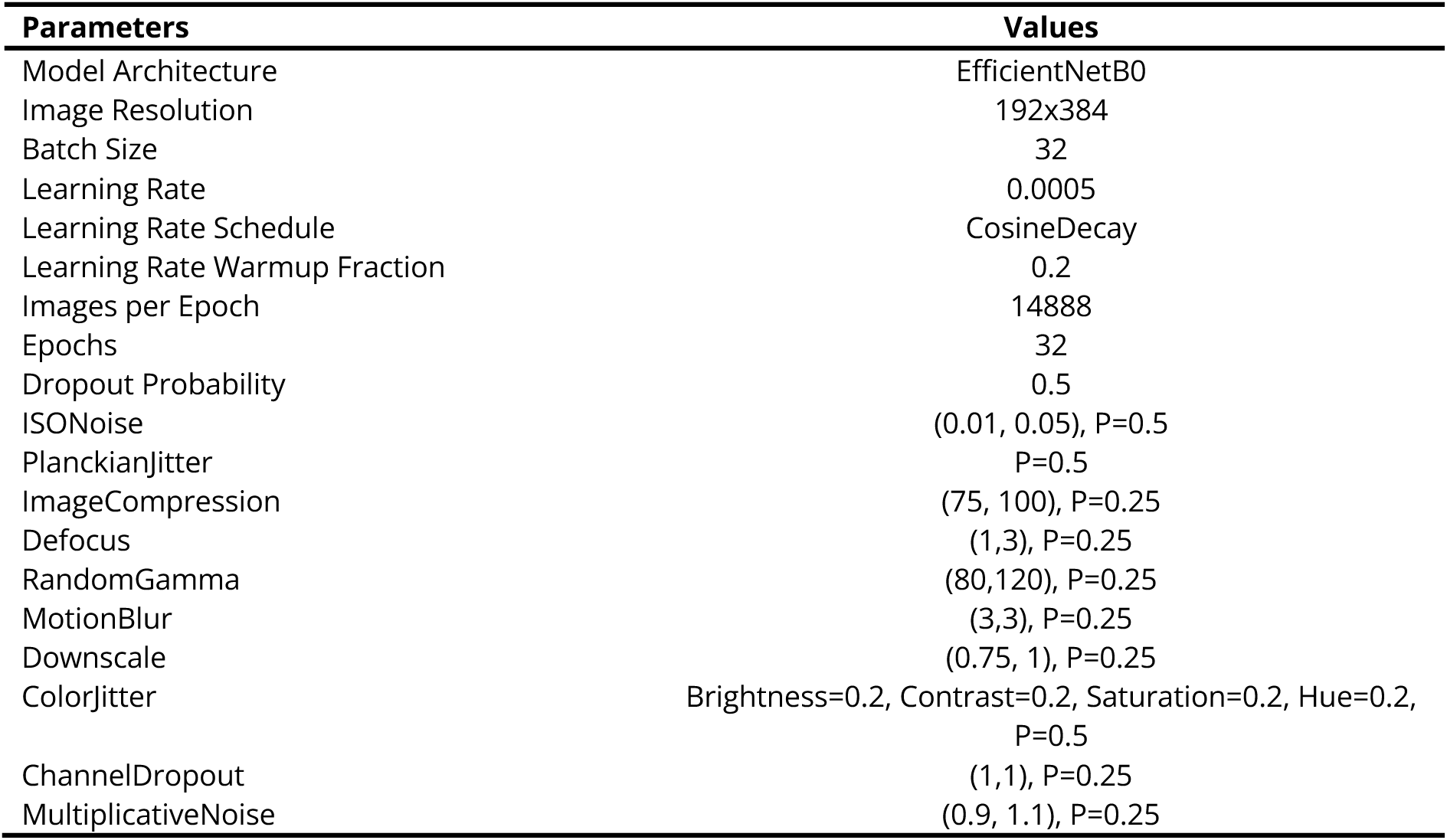

| Parameters | Values |
| --- | --- |
| Model Architecture | EfficientNetB0 |
| Image Resolution | 192x384 |
| Batch Size | 32 |
| Learning Rate | 0.0005 |
| Learning Rate Schedule | CosineDecay |
| Learning Rate Warmup Fraction | 0.2 |
| Images per Epoch | 14888 |
| Epochs | 32 |
| Dropout Probability | 0.5 |
| ISONoise | (0.01, 0.05), P=0.5 |
| PlanckianJitter | P=0.5 |
| ImageCompression | (75, 100), P=0.25 |
| Defocus | (1,3), P=0.25 |
| RandomGamma | (80,120), P=0.25 |
| MotionBlur | (3,3), P=0.25 |
| Downscale | (0.75, 1), P=0.25 |
| ColorJitter | Brightness=0.2, Contrast=0.2, Saturation=0.2, Hue=0.2, P=0.5 |
| ChannelDropout | (1,1), P=0.25 |
| MultiplicativeNoise | (0.9, 1.1), P=0.25 |

